# Collagen-VI expression is negatively mechanosensitive in pancreatic cancer cells and supports the metastatic niche

**DOI:** 10.1101/2022.03.03.482797

**Authors:** Vasileios Papalazarou, James Drew, Amelie Juin, Heather J. Spence, Colin Nixon, Manuel Salmeron-Sanchez, Laura M. Machesky

## Abstract

Pancreatic cancer is a deadly disease with high rates of metastasis, though how tumor cells establish metastatic lesions is not fully understood. A key feature of primary pancreatic tumors is extensive fibrosis due to deposition of extracellular matrix. While pancreatic cancer cells are programmed by stimuli derived from a stiff ECM, metastasis requires loss of attachment as well as adaptation to a softer microenvironment upon reaching distant sites. Growing evidence suggests that stiff ECM influences pancreatic cancer cell behaviour. Here we argue that this influence is reversible and that pancreatic cancer cells can be reprogrammed upon sensing of soft substrates. Through use of engineered polyacrylamide hydrogels with tuneable mechanical properties, we show that Collagen-VI is specifically upregulated on soft substrates, due to a lack of integrin engagement and low YAP1 activity. Collagen-VI supports migration *in vitro* and metastasis formation *in* vivo. Metastatic nodules formed by pancreatic cancer cells lacking *Col6a1* expression, were characterised by stromal cell-derived collagen-VI deposition, suggesting that collagen-VI, either cancer or stroma derived, is an essential component of the metastatic niche.

**Summary Statement:** Collagen-VI is expressed by pancreatic tumors and metastases in a mechanosensitive way to promote niche colonisation.

## Introduction

The dissemination of malignant cells from primary tumors to distant sites and the formation of metastatic nodules is a key step in cancer progression and aggressiveness. An essential component of the metastatic niche is the extracellular matrix (ECM) – the collection of extracellular proteins that provide the three-dimensional (3D) scaffold within which cells organise to form complex structures. It is emerging that cancer cell-ECM interactions have important roles in the establishment of metastatic tumors at all stages of the metastatic cascade (Drew and Machesky, 2021). These interactions depend on not only the biological components of the ECM but also its physical and mechanical properties.

The ability of cancer cells to respond to the physical properties of their surroundings is increasingly recognised as crucial to disease progression (Broders-Bondon et al., 2018). In solid tumors such as pancreatic ductal adenocarcinoma (PDAC), deposition of extracellular matrix (ECM) leads to extensive matricellular fibrosis that is linked to cancer aggressiveness (Mahadevan and Von Hoff, 2007; Perez et al., 2021) and contributes to its dismal 5-year survival rates (Siegel et al., 2021). Increased tissue stiffness has been shown to drive extravasation from the primary tumor through initiating epithelial-mesenchymal transition (EMT) in PDAC cells and stimulating invasion (Rice et al., 2017). However, cells escaping the primary tumor must survive and grow in low-adhesion and soft tissue environments such as the liver and lung to form metastatic lesions (Yachida and Iacobuzio-Donahue, 2009). Given that metastatic dissemination is the leading cause of death in pancreatic cancer (Balaban et al., 2017), understanding the adaptations that PDAC cells undergo during these transitions is of significant importance. Specifically, whether sensing of stiff ECM irreversibly modulates cancer cell behaviour or whether cancer cells can actively respond to low stiffness stimuli remains unclear.

Increased tissue stiffness and tension impacts cancer cell identity and behaviour through complex signalling pathways collectively referred to as “mechanosensation” (Papalazarou et al., 2018). The fundamental process of mechanosensation involves integrin receptor engagement of extracellular substrates, actin cytoskeleton remodelling and activation of key transcriptional regulators such as Yes-associated protein-1 (YAP1) (Panciera et al., 2017). PDAC cells use these mechanosensitive pathways to direct migration (Laklai et al., 2016), control metabolism (Papalazarou et al., 2020) and chemoresistance (Rice et al., 2017). Although some fundamental components of these pathways have been elucidated, much is still unknown about how mechanosensitive pathways are utilised by pancreatic cancer cells at different stages of disease progression.

In primary PDAC, the majority of ECM is deposited by stromal cells (in particular cancer-associated fibroblasts (CAFs)) that secrete large amount of fibrillar ECM components such as fibronectin and collagen-I, increasing tissue stiffness (Elyada et al., 2019). Unfortunately, approaches aimed at targeting ECM deposition in mouse models of PDAC (Ozdemir et al., 2014) and clinical trials (Doherty et al., 2018) have proven unsuccessful, likely due to the loss of ECM as a physical barrier to restrain the tumor (Bhattacharjee et al., 2021; Jiang et al., 2020). Therefore, a more nuanced understanding of the relationship between ECM deposition, tissue stiffness, and PDAC development is required.

Recent proteomic studies have revealed that alongside CAFs, cancer cells themselves produce a wide range of ECM proteins that contribute to the TME (Tian et al., 2019). Indeed, expression of cancer-cell derived ECM proteins correlate with poor survival in PDAC (Tian et al., 2019) suggesting they exert important roles in tumor progression. These functions may have relevance in metastatic dissemination, where cancer cells must quickly form a microenvironment to support their survival and growth (Drew and Machesky, 2021; Jiang et al., 2020). Indeed, a recent study showed that various PDAC-derived ECM components had roles in supporting metastatic dissemination (Tian et al., 2020); though our understanding of the functional roles of PDAC-derived ECM, and the mechanisms controlling its expression, remains limited. Given the intimate relationship between ECM and tissue stiffness, it is also unclear whether PDAC cells regulate their own ECM production in response to changes in their physical environment.

In this study, we define a link between substrate stiffness, PDAC ECM expression, and metastatic potential. By combining the use of mechanically tuneable hydrogels with RNA sequencing transcriptomic analyses, we observed that in environments of reduced stiffness pancreatic cancer cells upregulate the expression of genes involved in ECM production. This was prominently manifested by upregulation of three genes involved in Collagen-type VI synthesis: *Col6a1*, *Col6a2* and *Col6a3*. Focusing on Collagen-VI, we show that Collagen-VI upregulation is a response to a lack of integrin-based mechanosensation and diminished YAP nuclear localisation on soft substrates. Further, we find that PDAC cell-derived Collagen-VI supports invasive behaviour *in vitro* and metastatic potential of pancreatic cancer cells *in vivo*.

## Materials and methods

### Cell culture

All cell lines used in this study were cultured at 37°C under 5% CO_2_ in a humidified incubator and were tested regularly for mycoplasma contamination. Primary murine KPC PDAC and human PANC-1 cells were cultured in high-glucose DMEM supplemented with FCS (10%), glutamine (2mM), sodium pyruvate (0.11 g L^−1^), penicillin (10,000 units ml^−1^) and streptomycin (10,000 units ml^−1^). PDAC-A and PDAC-B cell lines were a gift from Jennifer Morton and Saadia Karim, and were isolated from the tumors of *Pdx1*-cre;LSL-*Kras*^G12D/+^;LSL-*Trp53*^R172H/+^ (KPC) mice either with a mixed or pure C57BL/J background.

### Reagents and Oligos

Human Fibronectin Protein (1918-FN-02M, R&D Systems), Fibronectin bovine plasma (F1141, Sigma-Aldrich), Concanavalin A from *Canavalia ensiformis* (L7647, Sigma-Aldrich), Poly-L-Lysine (P4707, Sigma-Aldrich), Aphidicolin from *Nigrospora spaherica* (A4487, Sigma-Aldrich), Corning Matrigel Basement Membrane Matrix (354234, Corning), Rat tail Collagen I (354249, Corning), Sigmacote (SL2, Sigma-Aldrich), sulfosuccinimidyl 6-(4’-azido-2’-nitrophenylamino)hexanoate (sulfo-SANPAH; 22589, Thermo Fisher Scientific), 3-(Acryloyloxy)propyltrimethoxysilane (L16400, Alfa Aesar), Acrylamide 40% Solution (A4058, Sigma-Aldrich), N,N’-methylene-bis-acrylamide 2% solution (M1533, Sigma-Aldrich), N,N,N’,N’-Tetramethylethylenediamine (TEMED; T9281, Sigma-Aldrich), Ammonium persulfate (A3678, Sigma-Aldrich), Ammonium hydroxide (221228, Sigma-Aldrich), Hydrogen peroxide (30%w/w; 31642, Sigma-Aldrich), Puromycin dihydrochloride (A1113803, Thermo Fisher Scientific), Lipofectamine RNAiMAX (13778150, Thermo Fisher Scientific), Lipofectamine 2000 (11668019, Thermo Fisher Scientific), Lullaby (FLL73000, OZ Biosciences), Amaxa Cell Line Nucleofector Kit V (VCA-1003, Lonza), Precision Red Advanced Protein Assay (ADV02-A; Cytoskeleton), calcein AM (C1430, Thermoo Fisher Scientific).

For CRISPR/Cas9-mediated genome editing, the following sequences were cloned into pSpCas9(BB)- 2A-Puro (PX459) V2.0 (gift from Feng Zhang, Addgene #62988) targeting mouse *Col6a1*: 5’- GTA CTT GAC CGC ATC CAC GC −3’(Col6a1^CRISPR#01^), 5’- TTG AGC TCA TCG CGG CCA C −3’ (Col6a1^CRISPR#02^), 5’- CTT GAT CGT GGT GAC CGA C −3’ (Col6a1^CRISPR#03^), 5’- ACT TGA TCG TGG TGA CCG A −3’ (Col6a1^CRISPR#04^) and mouse *Yap1*: 5’- ACT TGA TCG TGG TGA CCG A −3’ (YAP^CRISPR#01^). For siRNA mediated silencing the following sequences were used targeting mouse YAP, 5’- ACC AGG TCG TGC ACG TCC GC −3’ (YAP^siRNA#01^), 5’-ATGGAGAAGTTTACTACATAA-3’ (YAP^siRNA#02^) and mouse ILK, 5’- CCG CAG TGT AAT GAT CGA TGA −3’ (ILK^siRNA#01^) and 5’- CTC TAC AAT GTT CTA CAT GAA −3’ (ILK^siRNA#02^).

The following DNA constructs were used, pEFGP-C1 (host lab), pEGFPC1/GgVcl1-258 (VD1, a gift from Susan Craig, Addgene #46270 (Cohen et al., 2006)), EGFP-talin1 head (a gift from Anna Huttenlocher, Addgene plasmid #32856, (Simonson et al., 2006)). The L325R mutation was introduced to the EGFP-talin1 head construct using the Q5-site directed mutagenesis kit (NEB, E0554).

### Polyacrylamide hydrogel preparation

Polyacrylamide hydrogels were prepared at 0.7 kPa, 7 kPa and 38 kPa stiffness values as previously described (Papalazarou et al., 2020). Polyacrylamide hydrogels were treated with 0.2 mg ml^−1^ sulfo-SANPAH solution in MilliQ water (Thermo Fisher Scientific, 22589) followed by UV irradiation (365 nm) for 10 min. Hydrogels were extensively washed with 50 mM HEPES buffer (pH 8.5), incubated overnight with fibronectin (10 μg ml^−1^) and washed extensively in PBS before use.

### ECM coatings

Glass coverslips were washed with Ethanol, oven-dried and then coated with either fibronectin (10μg mL^−1^) for 60min at room temperature, Concanavalin A (ConA; 10 μg mL^−1^) or Poly-L-Lysine (PLL; 0.5 mg mL^−1^) for 16 hours at 4°C. Coated coverslips were washed in PBS and cells (2×10^4^ cells per mL) were plated and cultured for 16 hours in DMEM-10% FBS.

### RNA extraction, RNA sequencing and qRT-PCR

Total RNA extraction and purification from cells was performed using the RNeasy Mini Kit (Qiagen) combined with RNaseFree DNase (QIAGEN) treatment, according to the manufacturer’s instructions. Measurements of RNA concentration and purity were routinely performed using NanoDrop2000C (Thermo Fisher Scientific) before downstream processing. For RNA sequencing, RNA quality was tested using the Agilent Technologies 2200 TapeStation instrument according to the manufacturer’s instructions. Briefly, 5 μL of RNA sample buffer were mixed with 1 μL of RNA ladder and added to the first tube of a RNAse-free mini-tube strip. Then, 5 μL of RNA sample buffer were mixed with 1 μL of RNA sample and loaded to the strip. Samples were vortexed and centrifuged for 1 min at 2000 rpm. Samples were heated to 72°C for 3 min and place on ice for 2 min. Samples were centrifuged for 1 min at 2000 rpm and loaded into the Agilent 2200 TapeStation instrument.

Libraries were prepared using the Library Prep kit (Illumina TruSeq RNA Sample prep kit v2) and were run on an Illumina NextSeq500 platform using the High Output v2 75cycles (2×36cycle Paired End, single index) sequencing Kit. Quality checks on the raw RNASeq data files were done using FastQC (http://www.bioinformatics.bbsrc.ac.uk/projects/fastqc) and FASTQ screen (http://www.bioinformatics.babraham.ac.uk/projects/fastq_screen/) tools. RNASeq reads were aligned to the GRCm38 version of the mouse genome using tophat2 version 2.0.13 with Bowtie version 2.2.4.0.

Expression levels were determined and statistically analysed by a combination of HTSeq version 0.7.2 (http://www-huber.embl.de/users/anders/HTSeq/doc/overview.html), the R 3.3.3 environment, utilizing packages from the Bioconductor data analysis suite and differential gene expression analysis based on a generalized linear model using the DESeq2. Significantly changed genes (p_adj_<0.05) were submitted to DAVID for gene ontology (GO) analysis. KEGG pathway analysis was performed for genes demonstrating an increase (Up) or decrease (Down) in RNA expression between conditions. Significant KEGG GO Terms were identified (p_adj_<0.05). Hierarchical clustering of log_2_ Fold Changes in gene expression was performed on the basis of Euclidean Distance using complete linkage and visualised using the Rstudio, v.1.1.453 environment. We also performed DESeq analysis as well as Category netplot (CNET) analysis using the Bioconductor data analysis suite and the Rstudio, v.1.1.453 environment. The matrisome geneset source was taken from: https://www.gsea-msigdb.org/gsea/msigdb/cards/NABA_MATRISOME (accessed 02.2022) cDNA synthesis with either DyNAmo cDNA synthesis kit (F-470L, Thermo Fisher Scientific) or Maxima First Strand cDNA synthesis kit (K1641, Thermo Fisher Scientific). Then qRT-PCR was performed using the DyNAmo HS SYBR Green qPCR kit (F410L, Thermo Fisher Scientific). PCR was performed on a C1000 Thermal Cycler (CFX96 Real time system, BioRad) as follows: 3min at 95ºC, 40-cycles of 20s 95°C, 30s 57°C, 30s 72°C and final 5min 72°C. Relative mRNA quantification was performed using the 2^-ddCT method for multiple genes. Cdk2-Fw: TGAAATGCACCTAGTGTGTACC; Cdk2-Rv: TCCTTGTGATGCAGCCACTT; Col6a1-Fw: CGTGGAGAGAAGGGTTCCAG; Col6a1-Rv: GTCTCTCCCTTCATGCCGTC.

### Western blotting

Proteins were isolated in radioimmunoprecipitation (RIPA) lysis buffer (150mM NaCl, 10mM Tris-HCl pH7.5, 1mM ethylenediaminetetraacetic acid (EDTA), 1% Triton X-100, 0.1% SDS, 1X protease and phosphatase inhibitors (Roche)). Protein concentration was determined using Precision Red Advanced Protein Assay (Cytoskeleton, Inc.). 20ug protein were loaded onto 8-12% SDS-PAGE gels and transferred onto nitrocellulose membranes. Membranes were blocked (3%BSA / TBST) and incubated for 16h in 4°C with one of the following antibodies: anti-Collagen VI (1:1000; ab182744, Abcam), anti-YAP (1:1000; 14074, Cell Signaling Technology), anti-ILK (1:1000; 3856S, Cell Signaling Technology), anti-ERK1/2 (1:1000; 9102, Cell Signaling Technology), anti-GAPDH (1:1000; MAB374, Millipore) and anti-αTubulin (1:3000; T6199, Sigma). Protein detection was achieved using Alexa-Fluor conjugated secondary antibodies and signal was imaged using the LI-COR Odyssey CLx (LI-COR Biosciences) system. All images were analysed using Image Studio Lite software, version 5.2.5.

### Cell transfection and genetic modifications

Cells were typically transfected in suspension. 5 μg of DNA was used to transfect the cells using the Amaxa Cell Line Nucleofector kit (Lonza Bioscience) according to the manufacturer’s instructions. To generate stable cell lines, stably transfected cells were selected with G-418 (G418S, Formedium) followed by fluorescence activated cell sorting (FACS). For FACS, gating was performed by cell size, live/dead and GFP-positive signal.

For transfection of adhered cells, 1 μg of DNA was used with Lipofectamine 2000 reagent (Thermo Fisher Scientific) according to the manufacturer’s instructions. For siRNA-mediated YAP silencing, *Yap* targeting siRNA oligos (25nM; Qiagen) or control siRNA (25nM; AllStars Negative Control siRNA, Qiagen) were used to transfect the cells using the Lipofectamine RNAiMAX transfection reagent (Thermo Fisher Scientific) according to the manufacturer’s instructions. YAP-silenced cells were analysed 48h post-transfection. For siRNA-mediated ILK silencing, ILK targeting oligos (15nM; Qiagen) or control siRNA (15nM; AllStars Negative Control siRNA, Qiagen) were used with the Lullaby transfection reagent (OZ Biosciences) according to the manufacturer’s instructions. Two-step siRNA delivery with 48h interval was performed and ILK-silenced cells were analysed 24h post-transfection. For CRISPR/Cas9-mediated knockout, cells were transfected with 5μg of selected plasmid (control, empty vector (EV) or containing guide RNAs using Amaxa Cell Line Nucleofector kit (Lonza Bioscience) according to the manufacturer’s instructions. Stably transfected cells were selected using puromycin (2 μg mL^−1^).

### Immunofluorescence

Typically, 2×10^4^ cells per cm^2^ were seeded on 19-mm diameter coverslips or PAAm hydrogels for 16 hours. Cells were washed 1x in PBS and fixed in 4% PFA for 10 min. Then they were permeabilised with 0.1% Triton X-100 for 5 min followed by 30 min incubation in blocking buffer (1%BSA / PBS). Cells were incubated for 60 min with the following primary antibodies: anti-YAP (1:100; 14074, Cell Signaling Technology), anti-phosphoPaxillin (1:400; 2541, Cell Signaling Technology) and anti-Collagen VI (1:200; ab182744, Abcam). Detection was performed using the following secondary antibodies: Alexa Fluor 488 donkey anti rabbit (1:500 dilution; A21206, Invitrogen), Alexa Fluor 594 donkey anti mouse (1:500 dilution; A21203, Invitrogen). Nuclei were visualised with DAPI (0.5μg mL^−1^; D1306, Invitrogen) and F-actin with Alexa Fluor 647 Phalloidin (1:100 dilution; A22287, Invitrogen) incubation along with secondary antibodies. Coverslips were mounted using ProLong Diamond antifade reagent (P36965, Invitrogen).

Images were acquired using a Zeiss 880 Laser Scanning Microscope with Airyscan equipped with a Plan-Apochromat 63x/1.4 oil DIC M27 objective) or a Zeiss 710 confocal microscope equipped with Ec Plan Neofluar 40x/1.30 and Plan-apochromat 63x/1.40 oil objectives. All images were processed with Fiji software (ImageJ v2.0.0).

Focal adhesion analysis on Fig.3B was performed using the Fiji software (ImageJ v2.0.0) and the Analyze particles function applying a minimum size threshold of 0.25μm^2^. Cell shape analysis on Fig.S4A was performed by manually drawing around the cell perimeter based on F-actin staining and measuring the area and the shape descriptors using the Fiji software (ImageJ v2.0.0). Cell shape analysis on Fig.S4C was performed using Cell Profiler software (v3.0.0; CellProfiler), applying a mask for cell area based on F-actin staining and a mask for nuclei. YAP nuclear to cytosolic ratios on Fig.S1B were calculated using the Fiji software (ImageJ v2.0.0) to quantify mean YAP fluorescence intensity on similar rectangular areas over and adjacent to the nucleus.

### Immunohistochemistry (IHC)

All Haematoxylin & Eosin (H&E), Immunohistochemistry (IHC) and Picro-Sirius red (PSR) staining was performed on 4μm formalin fixed paraffin embedded sections (FFPE) which had previously been oven baked at 60⁰C for 2 hours.

The following antibodies were stained on an Agilent AutostainerLink48, αSMA (A2547, Sigma-Aldrich), Collagen VI (ab182744, Abcam), Fibronectin (A0245, Agilent) and p53 CM-5 (P53-CM5P-L). FFPE sections were loaded into an Agilent pre-treatment module to be dewaxed and undergo heat induced epitope retrieval (HIER) either using low or high pH target retrieval solution (TRS) (K8005, K8004, Agilent). Sections for αSMA staining underwent antigen retrieval using low pH TRS. Sections for Collagen VI, Fibronectin and p53 staining underwent antigen retrieval using high pH TRS. All sections were heated to 97⁰C for 20 minutes in the appropriate TRS. After HIER all sections were rinsed in flex wash buffer (K8007, Agilent) prior to being loaded onto the autostainer. The sections then underwent peroxidase blocking (S2023, Agilent) for 5 minutes, then washed with flex wash buffer. Sections for αSMA were blocked using mouse-on-mouse kit (MKB2213-1, Vector Lab) before primary antibody application for 35 minutes at a previously optimised dilution (αSMA, 1/25000; Collagen VI, 1/1000; Fibronectin, 1/750; p53, 1/250). The sections were then washed with flex wash buffer before application of appropriate secondary antibody for 30 minutes. Sections for αSMA had Mouse Envision applied (K4001, Agilent) and Collagen VI, Fibronectin, and p53 sections had Rabbit Envision (K4003, Agilent) applied for 30 minutes. Sections were rinsed with flex wash buffer before applying Liquid DAB (K3468, Agilent) for 10 minutes. The sections were then washed in water, counterstained with haematoxylin z (RBA-4201-001 CellPath).

The following antibodies were stained on a Leica Bond Rx autostainer, F4/80 (ab6640, Abcam, and Ly6G (BE0075-1, BioXcell). All FFPE sections underwent on-board dewaxing (AR9222, Leica) and antigen retrieval using appropriate retrieval solution. Sections for F4/80 staining were retrieved using enzyme 1 solution (AR9551, Leica) for 10 minutes at 37°C. Sections for Ly6G underwent antigen retrieval using ER2 solution (AR9640, Leica) for 20 minutes at 95°C. Sections were rinsed with Leica wash buffer (AR9590, Leica) before peroxidase block was performed using an Intense R kit (DS9263, Leica) for 5 minutes. Sections were rinsed with wash buffer before all sections had the blocking solution applied from the Rat ImmPRESS kit (MP7444-15, Vector Labs) for 20 minutes. Sections were rinsed with wash buffer (AR9590, Leica) and then primary antibody applied at an optimal dilution (F4/80, 1/100; Ly6G, 1/60000). The sections were rinsed with wash buffer and had Rat ImmPRESS secondary antibody applied for 30 minutes. The sections were rinsed with wash buffer and visualised using DAB in the Intense R kit.

H&E staining was performed on a Leica autostainer (ST5020). Sections were dewaxed, taken through graded alcohols and then stained with Haem Z (CellPath, UK) for 13 mins. Sections were washed in water, differentiated in 1% acid alcohol, washed and the nuclei blu’d in scotts tap water substitute (in-house). After washing sections were placed in Putt’s Eosin (in-house) for 3 minutes.

Staining for PSR was performed manually on FFPE sections that were dewaxed and rehydrated through xylene and a graded ethanol series before washing in water. Rehydrated slides were stained for 2 hours in PSR staining solution (equal volumes of 0.1% Direct red 80 (Sigma Aldrich) and 0.1% Fast green (Raymond A Lamb) (both in distilled water) combined in a 1:9 dilution with aqueous Picric acid solution (VWR).

To complete H&E, IHC and PSR staining sections were rinsed in tap water, dehydrated through graded ethanol’s and placed in xylene. The stained sections were coverslipped in xylene using DPX mountant (SEA-1300-00A, CellPath).

Slides were scanned with a Leica Aperio AT2 slide scanner at 20x magnification and all analyses were performed with HALO software (Indica labs).

### Random migration assay

Cells were plated at 20,000 cells per cm^2^ onto fibronectin coated dishes and imaged for 16h using a Nikon TE2000 microscope with a Plan Fluor 10x/0.30 objective and equipped with a heated CO_2_ chamber. Images were analysed with Fiji software (ImageJ v2.0.0).

### Inverted invasion assay

100 μL of 50% Matrigel in PBS solution were allowed to polymerise per transwell for 30 min at 37°C. Cells were trypsinized, resuspended in medium, counted and 4×10^4^ cells were plated on the underside of the transwell filter. Transwells were then placed inside 24-well plates in a way that the cell suspension droplets are in contact with the base of the 24-well plates and were incubated for 2 hours at 37°C allowing cell attachment to the bottom side of the filter. Transwells were then washed 3x with 1 mL of serum-free medium. Chemotactic gradients were created by filling the upper chambers with medium containing 10% FBS and the bottom chambers were kept in FBS-free medium. Cells were allowed to invade the Matrigel plug for 3.5 days and stained for 60 min at 37°C with 4 μM of calcein AM. Serial optical sections at 15 μM interval were obtained using an Olympus FV1000 confocal microscope equipped with a UplanSApo 20x/0.74 objective. Images were analysed using Fiji software (ImageJ v2.0.0).

### Wound invasion assay

96-well Incucyte Imagelock plates were coated for 16h with Matrigel (100μg mL^−1^) at 37°C. PDAC cell monolayers were generated by plating 7×10^5^ cells per mL for 4h. Monolayers were wounded using the 96-pin WoundMaker (Essen Bioscience) and embedded in Matrigel (2.15 mg mL^−1^, diluted in DMEM-10%FBS). Following 60 min incubation at 37°C, fully supplemented media were added to each well and images were acquired at 60 min intervals by the IncuCyte Zoom system (Essen Bioscience). Wound recovery was analysed with the automated Incucyte Zoom software (Essen Bioscience) providing relative wound density over time.

### Intraperitoneal and intrasplenic transplantation assays and KPC mice

A list of mice used for IHC staining is provided in Supplementary Table 1. Mice were maintained in the Biological Services Unit of the Beatson Institute according to UK Home Office regulations and in compliance with EU Directive 2010/63 and the UK Animals (Scientific Procedures) Act 1986. All protocols and experiments were previously approved by the Animal Welfare and Ethical Review Body (AWERB) of the University of Glasgow and were accompanied by a UK Home Office project licence to J. Morton (70/8375, intrasplenic) and to L. Machesky PE494BE48 (intraperitoneal and KPC mice). Intraperitoneal (IP) transplantation assay of PDAC cells was performed as previously described (Juin et al., 2019). PDAC cells (1×10^6^) were resuspended in 100 μL PBS and introduced into each nude mouse (CD-1nu, females, 6-week old; Charles River Laboratories, Wilmington, MA) by intraperitoneal injection. Tumor nodules were quantified in the mesenterium and the diaphragm following 14 days from injection.

Instrasplenic assay was performed as previously described (Papalazarou et al., 2020). Following anaesthesia and transverse incision exposing the spleen, PDAC cells (1×10^6^) in 100 μl PBS were injected in the spleen of nude mice (CD-1nu, females, 6-week old; Charles River Laboratories, Wilmington, MA). Surgical clips were typically removed 7 days after surgery and mice were sacrificed 21 days after inoculation. Liver tumour burden was calculated as the percentage of tumour-bearing liver lobes over the total number of liver lobes.

### Statistics and reproducibility

All datasets were analysed and plotted using Prism 8 (v8.2.0; GraphPad Software) unless otherwise stated. Differences between groups were tested for normal distribution and analysed using the appropriate statistical test, as mentioned in each figure legend. Error bars represent SD, unless otherwise stated.

#### Acknowledgements

We acknowledge CRUK Beatson Institute Core Services and Advanced Technologies (C596/A17196), CRUK Core grant A24452 to LMM, CRUK Centre Studentship A18076 to LMM, MSS and VP, EPSRC Programme Funding to MSS (EP/P001114/1).We especially thank Ann Hedley, Peter Bailey, and Billy Clark (CRUK Beatson Institute, Glasgow) for their assistance with preparation and analysis of RNA sequencing experiments; Saadia Karim and Jennifer P. Morton (CRUK Beatson Institute, Glasgow) for providing the two KPC-derived cells lines (PDAC-A and PDAC-B) used throughout; Jennifer P. Morton for advice about intrasplenic transplants. T. Hamilton, C. Baxter and E. Onwubiko (CRUK Beatson Institute, Glasgow) for helping with intrasplenic surgery. Feng Zhang (Massachusetts Institute of Technology, Cambridge), Anna Huttenlocher (Harvard Medical School, Cambridge) and Susan Craig (Johns Hopkins School of Medicine, Baltimore) for provision and advice on use of plasmids.

## Results

### Matrix stiffness alters pancreatic cancer cell expression of extracellular matrix proteins

To understand how pancreatic cancer cells respond to substrate stiffness, we cultured two independent murine KPC cell lines (pancreas tumor cells cultured from *Pdx1*-cre;LSL −*Kras*^G12D^; LSL-*Trp53*^R172H^ mice (Morton et al., 2010)) on fibronectin-coated polyacrylamide hydrogels of three defined stiffnesses, as reported previously (Papalazarou et al., 2020) and investigated gene expression profiles by performing RNA sequencing (Fig. 1A). We chose 0.7kPA, 7kPA, 38kPA to model soft, intermediate and stiff tissue contexts respectively (Butcher et al., 2009), and included a glass coverslip condition (~2-4 GPa) as a reference for typical cell culture conditions. As stiffness of healthy pancreas is typically < 1kPa and PDAC tissue can reach 7-10 kPa these stiffness values have physiological relevance to PDAC (Rice et al., 2017). Immunofluorescent (IF) imaging revealed clear morphological differences between different conditions, with cells on soft substrate appearing round and clustered compared to the elongated cells with visible protrusions and stress fibres seen on stiff substrate and glass (Fig.1B). Further, we observed a dramatic redistribution of the mechanosensitive transcriptional regulator YAP1 (Zanconato et al., 2019) from the nucleus to the cytoplasm on soft substrate (Fig.S1A-B), indicating that the system is indeed driving mechanosensitive pathways in PDAC cells.

**Figure 1.**
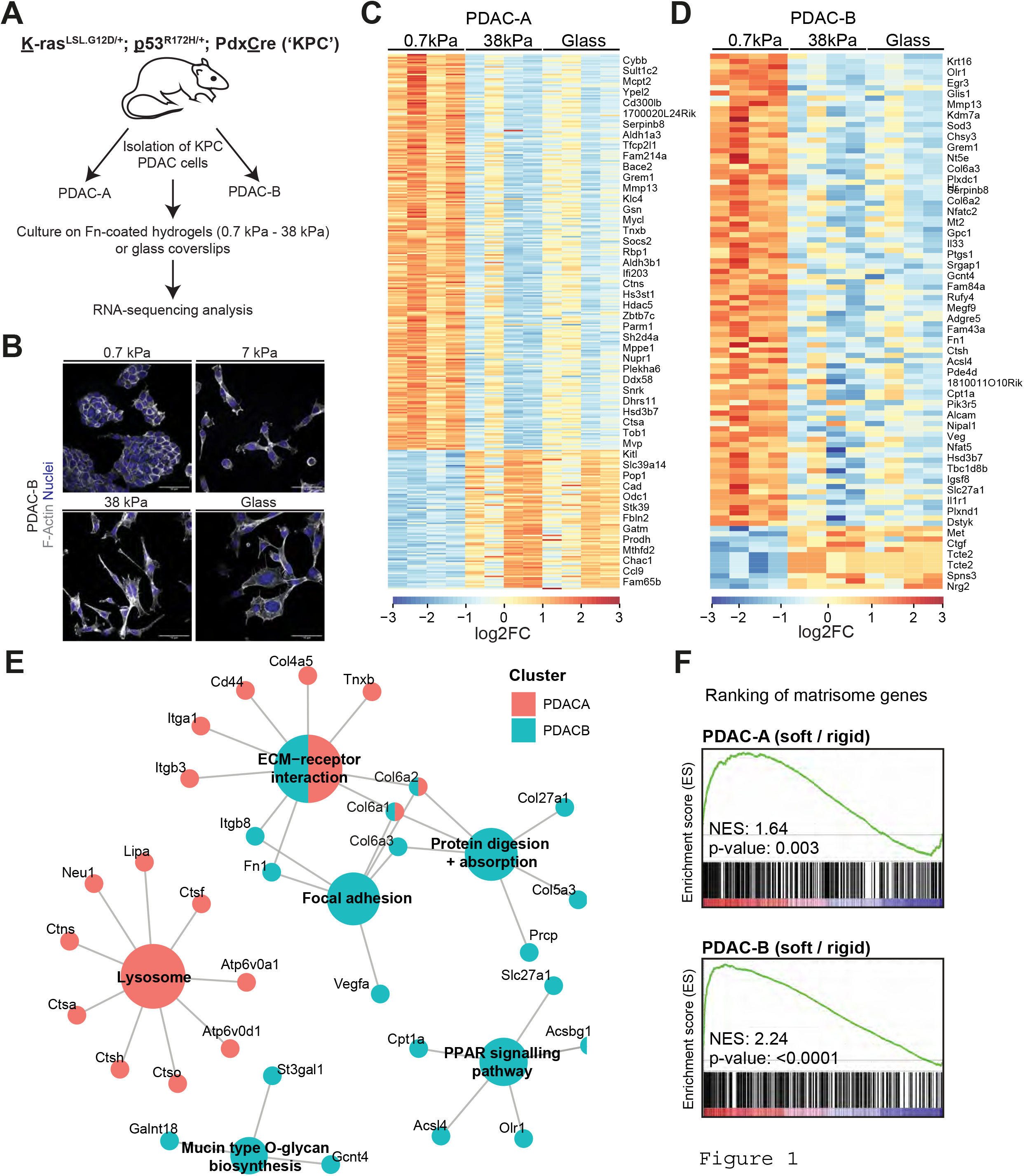
Low substrate stiffness promotes the expression of matrisome-related genes in PDAC cells. **A:** Schematic representation of RNA sequencing strategy of PDAC cells. Two KPC PDAC cell lines isolated from the *Kras*^LSL.G12D/+^;*Trp*53^R172H/+^; *Pdx*-Cre (‘KPC’) mouse model were cultured atop of 0.7 kPa, 7 kPa or 38 kPa Fibronectin (Fn)-coated polyacrylamide hydrogels or Fn coated glass coverslips for 24 hours. Total RNA was extracted and RNA sequencing was performed. **B:** Immunofluorescence of PDAC-B cells cultured atop of 0.7 kPa, 7 kPa or 38 kPa fibronectin-coated polyacrylamide hydrogels or fibronectin-coated glass coverships showing F-Actin (phalloidin, grey) or nuclei (Hoechst, blue). Scale bars, 50 μm. **C-D:** Expression heatmaps of differentially expressed genes of cells cultured on 0.7 kPa, 38 kPa hydrogels and glass coverslips. In PDAC-A cells **(C)**, 258 genes were upregulated and 92 downregulated at 0.7 kPa. In PDAC-B cells **(D)**, 90 genes were upregulated and 10 downregulated at 0.7 kPa (p_adj_ < 0.05; log2 fold change >1). In both cell lines, glass culture conditions show similarity in expression pattern with cells cultured on 38 kPa hydrogels. Data is from n = 4 independent replicates per condition for each cell line. **E:** CNET plot of KEGG pathway analysis based on upregulated genes in PDAC-A and B cells identified in **C-D**. Individual genes important to enrichment of these functional nodes are shown. Colours represent genes upregulated in either PDAC-A (red), PDAC-B (blue) or both (split). **F:** GWAS plot showing enrichment of matrisome geneset in ranked gene lists of both PDAC-A (top) and PDAC-B (bottom) cell lines.

To better understand the global changes in gene expression caused by different substrate stiffnesses, we conducted RNA sequencing of PDAC-A and PDAC-B cell lines on soft and stiff hydrogels, and glass. Hierarchical clustering of replicates revealed markedly distinct transcriptional profiles of the two cell lines regardless of ECM stiffness (Fig.S1C), but that within cell lines replicates from soft conditions tended to cluster separately (Fig.S1D). Indeed, analysis of differentially-expressed genes (DEGs) showed substantial changes in gene expression between soft substrate and both stiff hydrogels and glass conditions (Fig.1C-D). The observation that gene expression changes are consistent between stiff and glass conditions supports the idea that PDAC cells are mechanosensitive within a physiological range of substrate stiffnesses. Pathway analysis of genes upregulated on soft substrates revealed a surprising enrichment in genes associated with the ECM and substrate-interacting pathways (Fig.1E). Gene-level analysis of the ‘ECM-Receptor Interaction’ pathway showed a mix of ECM molecules (collagens, fibronectin) and integrins were being upregulated on soft substrates (Fig.S1E). To confirm this, a gene-set enrichment analysis between soft and stiff conditions was performed using the matrisome geneset from Naba et al. (Naba et al., 2012). Both cell lines showed a significant enrichment in matrisome gene expression on soft hydrogels (Fig.1F). Thus, PDAC cells show an orchestrated response to changes in mechanical stiffness of substrate characterised by increased expression of ECM-related genes.

### Collagen-VI is upregulated by pancreatic cancer cells on soft substrates downstream of ECM adhesion and YAP

Comparing the gene-level changes in PDAC-A and PDAC-B cell lines, we noticed that several subunits of Collagen-VI were upregulated on soft matrix in both lines (Fig.S1E). Collagen-VI is a fibrillar collagen that exists as a tetrameric assembly of α1 and α2 subunits, with possible inclusion of α-3/4/5/6 (Fig.2A). We first sought to confirm upregulation of Collagen-VI on soft substrates at the RNA and protein level. qPCR of *Col6a1* showed a gradual increase in mRNA levels across progressively softer substrates, which was significant when comparing the 0.7kPa hydrogel and glass conditions (Fig.2B). Increases in mRNA were shown to translate to protein expression via Western blots for Collagen-VI in both cell lines (Fig.2C-D). Again, a gradual increase in protein expression was observed across the substrate stiffnesses, highlighting the sensitivity of Collagen-VI expression in PDAC cells to physical properties of the substrate. IF of Collagen-VI in PDAC-A and a separate human PDAC cell line (PANC1) showed a clear increase in intracellular Collagen-VI staining as substrate stiffness was reduced (Fig.S2A-B). Finally, to assess whether Collagen-VI expression could be an adaptive response to slower proliferation levels of PDAC cells upon low substrate stiffness (Papalazarou et al., 2020), we treated cells with aphidicolin, an inhibitor of DNA replication (Fig.S3A). Upon aphidicolin treatment there was a trend for reduced Collagen VI expression, suggesting that the increase of Collagen-VI expression upon soft matrices is not a direct adaptation in response to lower proliferative capacity on soft matrix.

**Figure 2.**
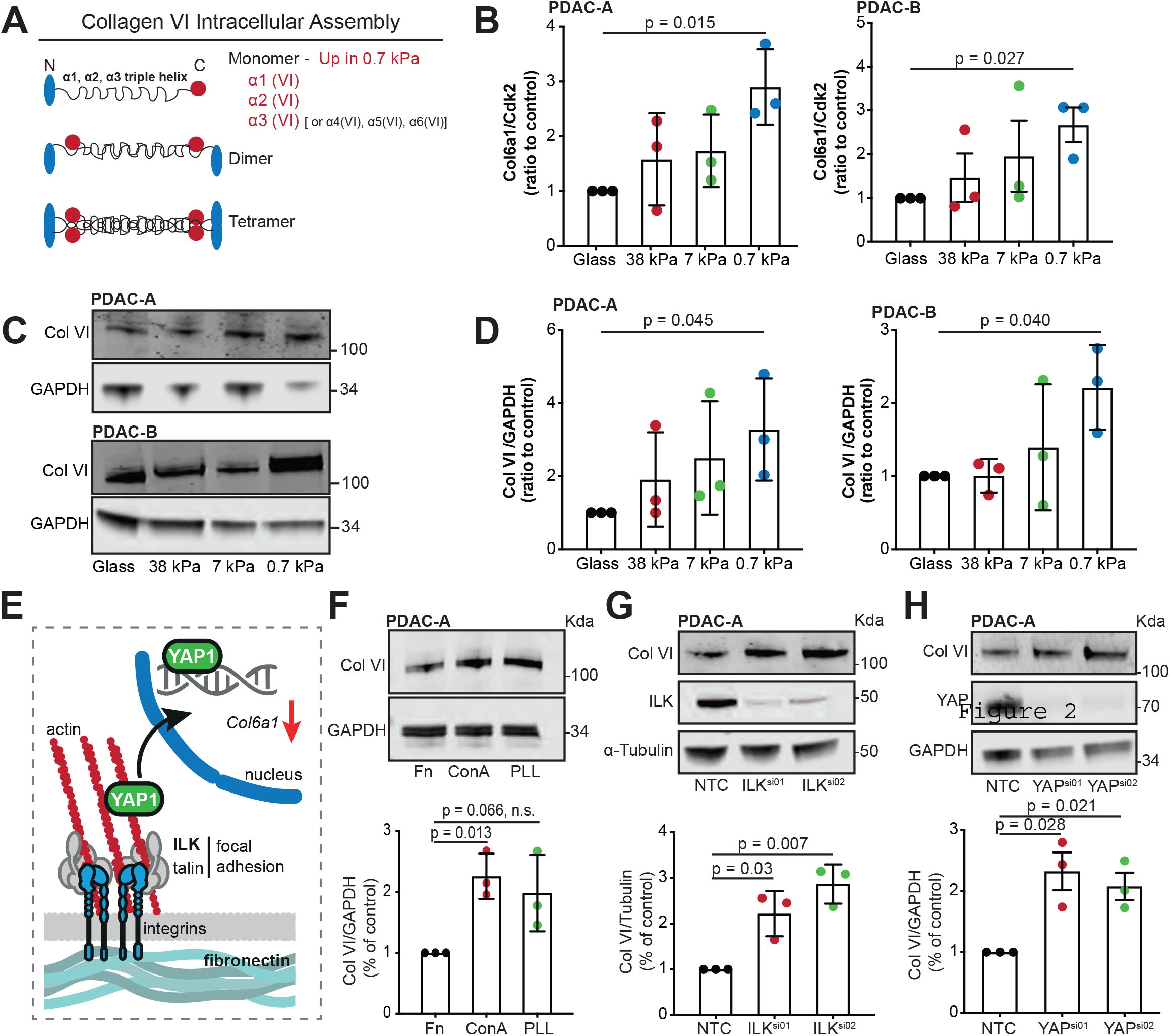
Collagen VI is upregulated in PDAC cells upon low substrate stiffness. **A:** Schematic representation of the collagen VI assembly process in cells. The collagen VI monomer is a triple helix encoded by 3 genes: *Col6a1*, *Col6a2* and *Col6a3*. In certain cases, the *Col6a3* chain can be replaced by *Col6a4* (not functional in humans), *Col6a5* or *Col6a6*. In B-D, PDAC-A and PDAC-B cells were cultured on 0.7, 7- and 38-kPa fibronectin-coated hydrogels and glass coverslips. **B:** Col6a1 mRNA expression was measured with qRT-PCR. Values are mean ± SD and relative to control expression (Cdk2) from 3 independent experiments. **C:** Collagen VI protein expression was measured by immunoblotting for Collagen VI (Col VI) and GAPDH (loading control). Blots are representative of three independent experiments. **D:** Densitometric quantification of protein in C. Values are mean ± s.d. of three independent experiments. **E:** Schematic representation of main adhesion-linked pathways involved in sensing and responding to substrate stiffness by pancreatic cancer cells. **F:** Top; PDAC-A cells were cultured on fibronectin (Fn, control), concanavalin A (ConA) or poly-l-lysine (PLL) coated glass coverslips and immunoblotting for Collagen VI and GAPDH (loading control) was performed. Bottom; Densitometric quantification of protein expression. Values are mean ± s.d. of three independent experiments. **G:** Top; Control (NTC) or ILK silenced (*Ilk*^si01^, *Ilk*^si02^) PDAC-A cells were immunoblotted for Collagen VI, ILK and α-Tubulin (loading control). Bottom; Densitometric quantification of protein expression. Values are mean ± s.d. of three independent experiments. **H:** Top; Control (NTC) or YAP silenced (YAP^si01^, YAP^si02^) PDAC-A cells were immunoblotted for Collagen VI, YAP and GAPDH (loading control). Bottom; Densitometric quantification of protein expression. All data in B-D and F-H are from 3 independent experiments. B, D, F, G, H: Statistical significance was assessed by two-tailed one-sample *t*-test on natural log-transformed values.

To better understand the mechanism driving increased expression of Collagen-VI on soft hydrogels, we conducted a range of experiments disrupting key steps in the integrin and YAP-linked mechanosensation pathway (Fig.2E). Firstly, to assess the requirement of integrin receptor engagement, PDAC-A cells were cultured on plastic coated with fibronectin, or two substrates known to allow cell attachment without engagement of integrin receptors, concanavalin A (conA) and poly-l-lysine (PLL) (Fig.2F). Collagen-VI expression was upregulated in both PDAC-A cells grown on conA and PLL, suggesting that loss of integrin receptor engagement drives Collagen-VI expression. Integrin-linked kinase (ILK) is a component of focal adhesion complexes that translate integrin engagement to intracellular signalling (Hannigan et al., 2005). Transfection of PDAC-A cells with two different siRNAs targeting *Ilk* led to 2-3 fold increase in collagen-VI expression (Fig.2G), that was also observed in PDAC-B cells (Fig.S3B). Overexpression of a dominant-negative Talin-head domain mutant that disrupts binding to integrins (L325R) also increased Collagen-VI expression compared to cells overexpressing a wild-type Talin-head domain or the vinculin head domain (VD1) in PDAC-A cells (Fig.S3C,D). Finally, we asked if Collagen-VI expression was directly altered by disruption of YAP, a major mechanosensitive transcriptional regulator that lies downstream of integrin-focal adhesion complex signalling. Silencing YAP expression with two siRNAs led to ~2-fold increase in Collagen-VI expression (Fig.2H). These results were confirmed using a CRISPR KO of *Yap1* in PDAC-A cells, additionally showing that Collagen-VI expression was higher in *Yap1* KO cells regardless of substrate stiffness (Fig.S3E-F). Taken together, these results show that upregulation of Collagen-VI expression on soft substrata is likely driven by loss of integrin receptor engagement and YAP activity.

### Cancer cell-derived collagen-VI supports migration invasive behaviours in PDAC cells

Prior studies have found a role for Collagen-VI in supporting cancer cell migration and invasion, though these studies did not investigate the influence of cancer cell-derived collagen-VI (Iyengar et al., 2005; Wishart et al., 2020). To ask if Collagen-VI altered pancreatic cancer cell morphology and migration, we cultured PDAC-A cells on either fibronectin, Collagen-VI or a 50:50 mix of the two (Fig.3A-B, S4A-B). Cells plated on Collagen-VI produced significantly less mature focal adhesions, indicating a less stable association with the substrate (Fig.3B). Supporting this, timelapse imaging over a 16-hour period revealed an approximate 2-fold increase in migration speed on collagen-VI (Fig.3B, S4B).

As these experiments utilise exogenous recombinant protein, it is unclear if cancer cell-derived Collagen-VI can support migratory behaviour in the same manner. To test this, we selected two of four CRISPR lines in PDAC-A cells in which a critical exon of *Col6a1* was deleted (Fig.3C). Cells appeared morphologically normal, though we did observe a slight increase in cell area in the Col6a1.03 line (Fig. 3D, S4C). We used two complementary experiments to investigate the invasive potential of these cells. Firstly, invasion into a Matrigel plug in response to a serum chemotactic gradient was used (Fig.3E). Analysis of the depth that cells invaded into the plug after 72 hours revealed a reduction in both CRISPR KO lines that was significant for Col6a1.04 (Fig.3F-G, S4D). Secondly, we utilised a 3D wound healing assay in which a Matrigel-embedded cell monolayer invades into wounded ECM (Fig.3H). We again observed a reduction in wound closure over a 50-hour period that was significant for the Col6a1.04 line (Fig.3I-K). Together, these results indicate that cell-derived Collagen-VI in pancreatic cancer cells supports invasive behaviours in a range of contexts.

**Figure 3.**
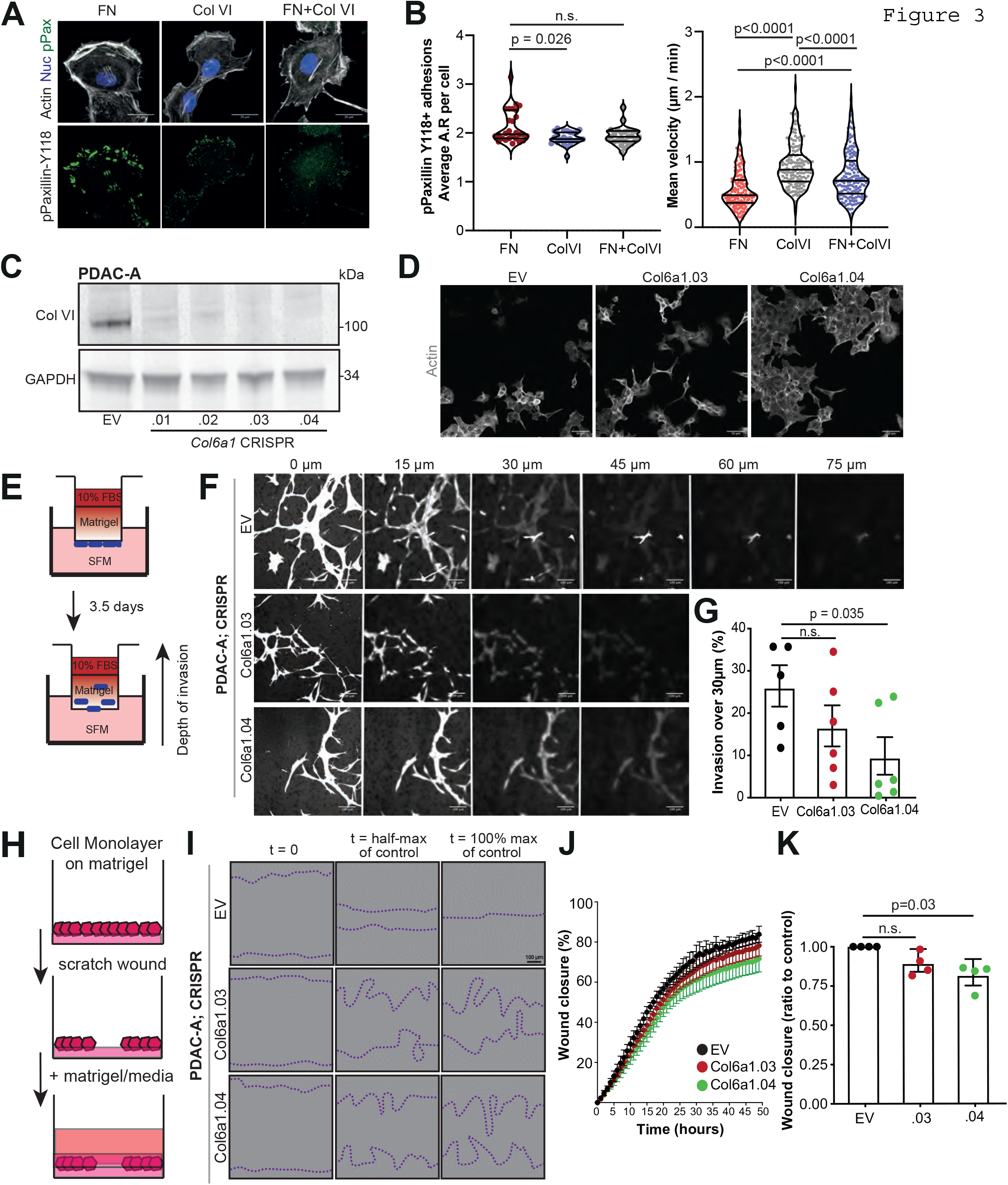
Collagen VI ECM supports migratory behaviour of PDAC cells *in vitro* and loss of Col6a1 expression delays invasion through recombinant basement membrane ECM. **A**: Top; immunofluorescence of PDAC-A cells cultured on fibronectin (FN), collagen VI (ColVI) or fibronectin and collagen VI (FN+ColVI) glass coverslips showing F-Actin (grey), phospho-Paxillin^Y118^ (green) and nuclei (blue). Bottom; Individual phospho-Paxillin^Y118^ channel (green). Scale bars, 20μm. **B**: Left; Quantification of p-Paxillin-positive particles from A showing average aspect ratio of focal adhesions per cell. Values are from *n* = 27 FN; *n* = 24, ColVI; *n* = 25, Fn+ColVI cells. Cells are from three independent experiments. Statistical significance was assessed by Kruskal-Wallis with Dunn’s multiple comparisons test. Right; Cell speed from Fig. S4B. Values are from *n* = 155 FN cells; *n* = 163, ColVI; *n* = 166, Fn+ColVI cells. Cells are from three independent experiments. Statistical significance was assessed by Kruskal-Wallis with Dunn’s multiple comparisons test. **C**: Control (EV) or Collagen VI depleted (Col6a1.01 – .04) mouse PDAC-A cell lines were immunoblotted for Collagen VI and GAPDH (loading control). Pictures are representative of 3 independent experiments. **D**: Immunofluorescence of control (EV) or Collagen VI depleted (Col6a1.03 and Col6a1.04) PDAC-A cell lines showing Actin (grey). Scale bars, 50μm. **E**: Schematic representation of the inverted invasion assay setup. **F**: Representative pictures from z-stack acquisitions of control (EV) or Collagen VI depleted (Col6a1.03 and Col6a1.04) PDAC-A cell lines invading Matrigel plugs showing Calcein staining after 72 hours of invasion. Scale bars, 100μm. **G**: Invasion over 30μm from F. Values are mean ± s.e.m. from 3 independent experiments. Statistical significance assessed with two-way Welch’s t-test. **H**: Schematic representation of the 3D ECM wound invasion assay setup. **I**: Representative pictures of control (EV) or Collagen VI depleted (Col6a1.03 and Col6a1.04) PDAC-A cells invading 3D ECM. Scale bar, 100 μm. **J**: Wound closure over time of control (EV) or Collagen VI depleted (Col6a1.03 and Col6a1.04) PDAC-A cells invading 3D ECM. Values are mean ± SD from 4 independent experiments. **K**: Relative wound closure of J normalised at t_1/2_ wound closure of control. Values are mean ± SD from 4 independent experiments. Statistical significance assessed by one-sample t-test on LN transformed values.

### Collagen-VI deposition increases during PDAC progression and is associated with poor survival

Having shown that pancreatic cancer cells display mechanosensitive expression of Collagen-VI, and that Collagen-VI supports their motility in 2D and 3D settings, we investigated whether cancer cell-derived Collagen-VI supported pancreatic cancer progression *in vivo*. Collagen-VI has well documented roles in certain cancers including breast (Wishart et al., 2020), ovarian (Sherman-Baust et al., 2003) and lung (Voiles et al., 2014). We began by investigating whether Collagen-VI expression is altered in the KPC mouse model of PDAC. We stained pancreata of normal (*Pdx1*-Cre^+/-^:*Kras*^+/+^:*Trp53*^+/+^) or KPC (*Pdx1*-Cre^+/-^:*Kras*^G12D/+^:*Trp53*^R172H/+^) mice at different stages of PDAC progression from PanIN I-III (10- and 15-week) to endpoint PDAC (Fig.4A). We noticed an upregulation of Collagen-VI expression with PDAC progression (Fig.4B). Further, Collagen-VI was mainly localised in the stroma, suggesting that stromal cells (particularly cancer associated fibroblasts) may be the primary depositors of this type of Collagen-VI in primary PDAC tissue, in agreement with a recent proteomic study (Tian et al., 2019). As primary PDAC tissue is typically stiff, it is unsurprising that cancer cells are not the major source of Collagen-VI in primary tissue. Indeed, Collagen-VI expression in liver metastasis samples of KPC mice (where cells will be experiencing softer substrate) showed markedly stronger staining that also appeared more cytoplasmic and PDAC cell-derived (Fig.4C).

**Figure 4.**
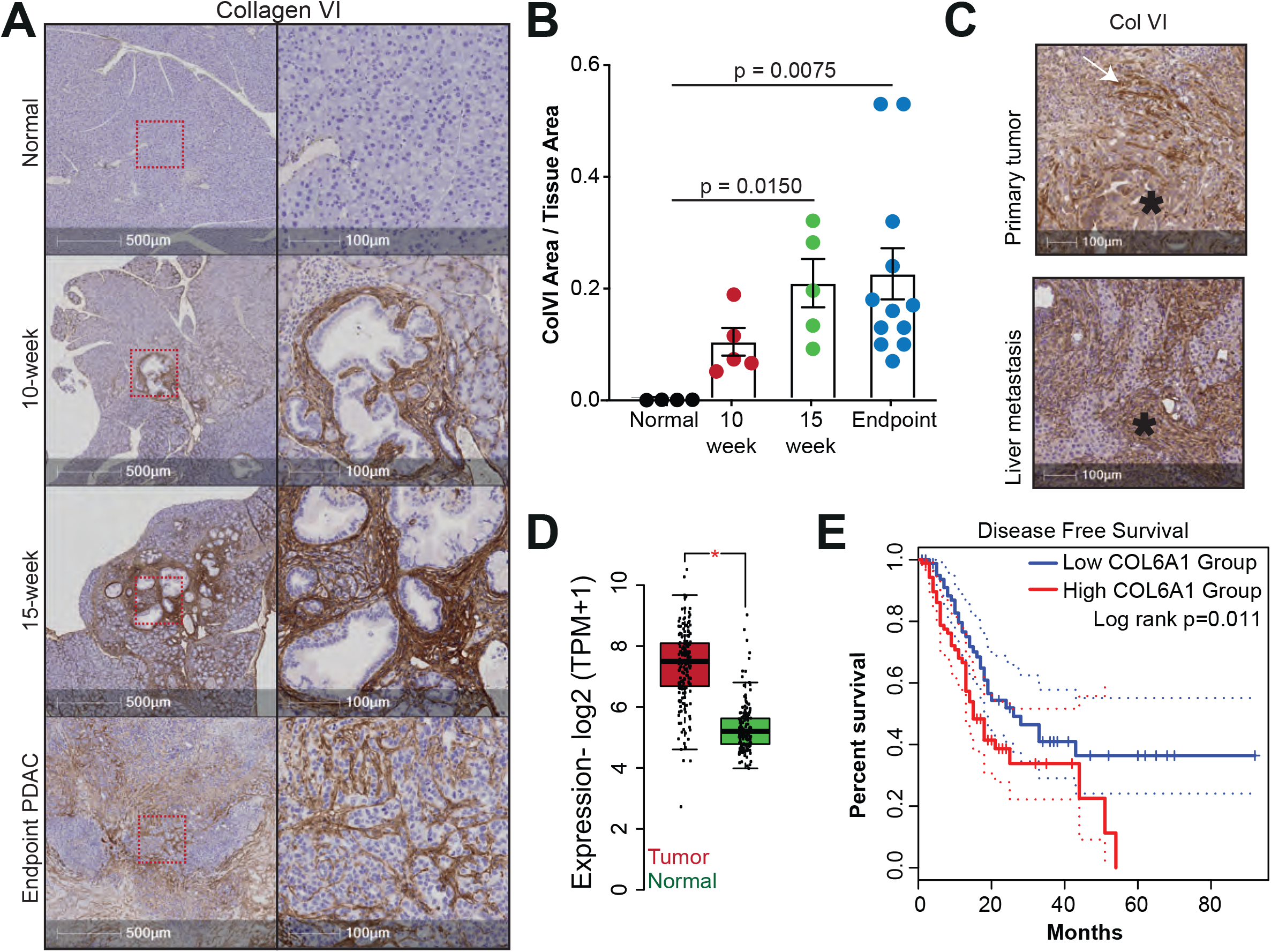
Collagen VI is upregulated during PDAC progression and correlates with reduction in disease-free survival. **A**: Immunohistochemistry of normal mouse pancreas (top panel), 10-week, 15-week and endpoint PDAC pancreas from KPC mice showing Collagen VI expression. Red boxes on left panel indicate magnified area (right panel). Scale bars, 500 μm (left) and 100 μm (right). **B**: Quantification of Collagen VI positive area per tissue area from (A). Values are mean ± s.e.m. from n = 4 normal, n = 5 10-week, n = 5 15-week and n = 12 mice. Statistical significance assessed by Kruskal-Wallis with Dunn’s multiple comparisons test. **C**: Representative images of immunohistochemistry from endpoint PDAC pancreas (primary tumor) and metastatic nodules in the liver from KPC mice showing Collagen VI expression. Scale bars, 100 μm. Asterisks indicate pancreatic tumor cells and are representative from n = 3 mice with matched primary tumor and liver metastasis samples. **D**: Expression of COL6A1 is upregulated in the tumors of pancreatic cancer patients compared to normal pancreas tissue specimens. Data taken from TCGA/GTEx (Tumor: n=179 patients; Normal: n=171 patients). Statistical significance assessed by two-sample t-test * p<= 0.05. **E**: Kaplan-Meier plot of disease-free survival stratifying patients based on *COL6A1* expression in pancreatic tumors. High *COL6A1* expression is associated with a significant reduction in disease free survival (Log rank p < 0.012). n = 89 patients with high *COL6A1* expression and n = 89 patients with low COL6A1 expression.

To confirm that these results were representative of human disease, we consulted the TCGA database of primary human PDAC tissue (Fig.4D-E). RNA levels of *COL6A1* were significantly increased in primary tumor vs normal pancreas samples (Fig.4D). Furthermore, high *COL6A1* expression was associated with reduced disease-free survival in these patients (Fig.4E).

### Collagen-VI expression supports establishment of pancreatic metastasis *in vivo*

We hypothesised that PDAC cells may upregulate Collagen-VI upon extravasation in the liver, contributing to the establishment of the metastatic niche. To test this hypothesis, we initially performed intraperitoneal injection (IP) experiments using a KPC mouse-derived cell line (KPC #127445) that we have used previously for such experiments (Juin et al., 2019) (Fig.5A). Two *Col6a1* CRISPR KO lines were used alongside an empty vector (EV) control to generate abdominal cavity and diaphragm tumors (Fig.5A, S5A). Interestingly, both *Col6a1* KO lines showed a trend for decreased tumor formation on the diaphragm (Fig.5B) and the peritoneum (Fig.5C) that was significant for the Col6a1.04 line, despite no changes in mouse weights (Fig.S5B). Histological analysis of these tumors confirmed the high levels of Collagen-VI expression in abdominal cavity tumors from EV cells and the significant loss of Col6a1 staining in both KO lines (Fig.5D). What Collagen-VI staining remained in *Col6a1* KO tumors appeared to colocalise with αSMA+ cells, which likely represent myofibroblast-like resident cells (Fig.S5C).

**Figure 5.**
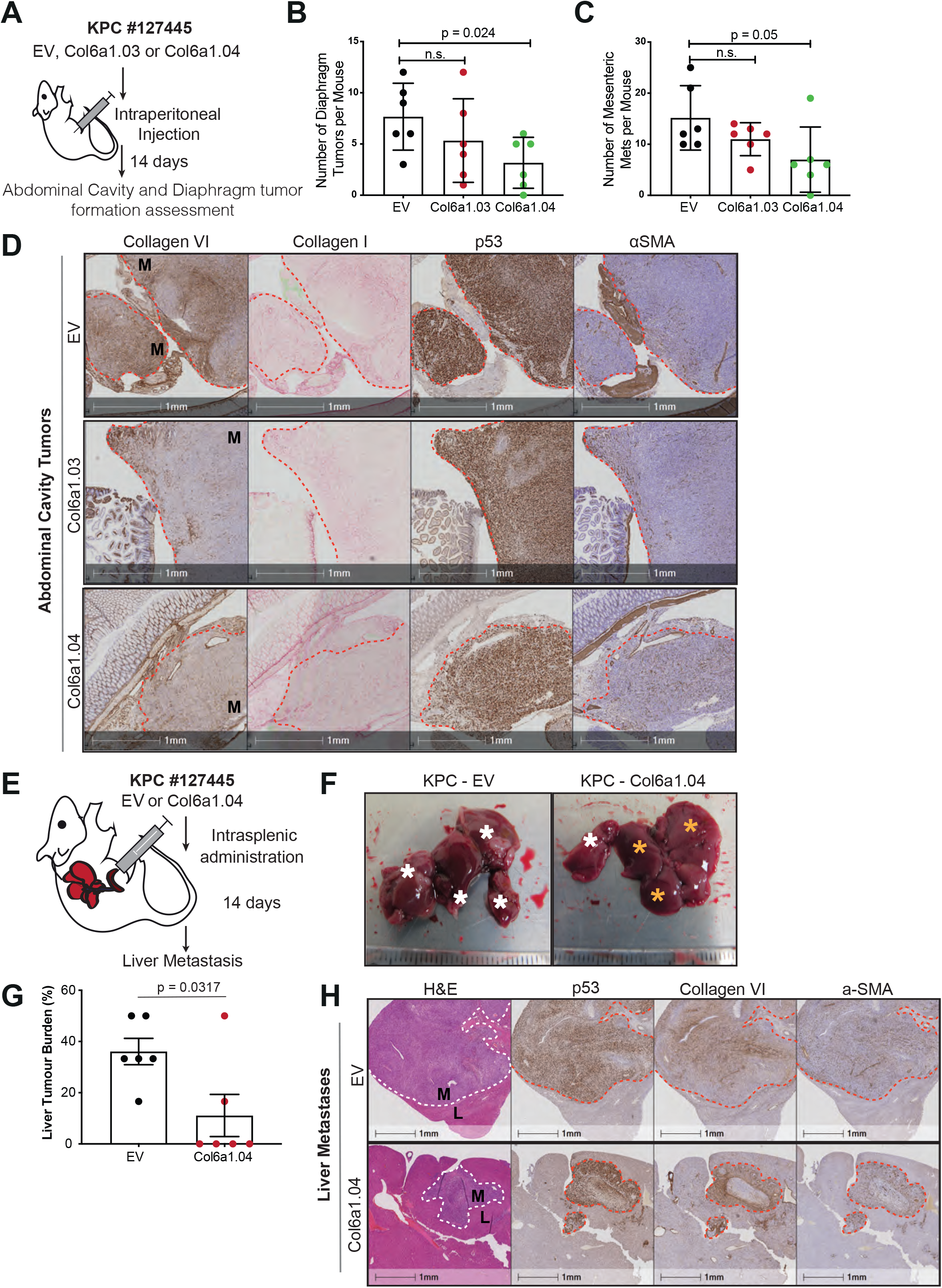
Collagen VI expression supports establishment of pancreatic metastasis in vivo. **A**: Schematic of intraperitoneal injection model. Control (EV) or Collagen VI depleted (Col6a1.03 and Col6a1.04) KPC cells were injected in the intraperitoneal cavity of nude (Cd1-nu) mice. After 14 days the mice were sacrificed, and the number of tumors was quantified before fixation of isolated tissues. **B:** Quantification of diaphragm tumors per mouse as indicated. Values are mean ± SD from n=6 EV, n=6 Col6a1.03 and n=6 Col6a1.04 mice. Statistical significance was assessed with Welch’s t-test. **C**: Quantification of abdominal cavity tumors per mouse as indicated. Values are mean ± s.d. from mice receiving control or CRISPR cells n=6 EV, n=6 Col6a1.03 and n=6 Col6a1.04 mice. Statistical significance assessed with Welch’s t-test. **D**: Representative immunohistochemistry images showing Collagen VI, Collagen I, p53 (marks tumor, with mutant p53 accumulation) and αSMA (marks fibroblasts) expression in abdominal cavity tumor nodules formed by intraperitoneal injection of control (EV) or Collagen VI depleted cells (Col6a1.03 and Col6a1.04). Red line denotes boundary between normal tissue and metastases (M). Scale bars, 1mm. E: Schematic of intrasplenic injection model. Control (EV) or Collagen VI depleted (Col6a1.04) KPC cells were transplanted into the spleen of nude (CD1-nu) mice and 14 days later formation of metastatic tumors in the liver was assessed. **F**: Representative liver pictures at the time of dissection. White asterisks indicate lobes with metastatic tumors and yellow asterisks indicate tumor-free liver areas. **G**: Liver tumor burden (% of liver lobes affected) of animals treated as indicated. Values are mean ± s.e.m. from n = 6 EV and n = 6 Col6a1.04 mice. Statistical significance was assessed with Welch’s t-test. **H**: Representative immunohistochemistry images showing H&E, p53, Collagen VI and α-SMA expression in liver sections from F as indicated. White or red dotted line denotes boundary between normal liver (L) tissue and metastases (M). Scale bars, 1mm.

The liver is a common site of metastasis in pancreatic cancer. To further validate these observations and since IP injections do not represent the metastatic cascade, we decided to assess the role of PDAC-derived Collagen-VI in formation of liver metastases using the model of intrasplenic (IS) transplantations (Fig.5E). Interestingly, Collagen-VI KO cells (Col6a1.04) displayed a reduced ability to form metastases in the liver compared with control cells (EV) following IS transplantation (Fig.5F-G). Histological staining again showed that Collagen-VI was deposited primarily in areas with high αSMA+ cells in KO conditions and was more diffuse in EV tumors (Fig.5H). Further, KO metastases typically had strong Collagen-VI deposition surrounding tumor nodules, likely of stromal origin, suggesting that presence of Collagen-VI is required to support metastasis formation - regardless of its source. We validated these histological results by generating liver metastases with PDAC-B CRISPR control (EV) or Col6a1 KO (Col6a1.03) cell lines (Fig.S5D-E). Analysis of common ECM markers and stromal cell types showed no significant differences between EV and Col6a1.03 conditions, though there was a trend for 1.5-2-fold more abundance of fibroblasts (αSMA+), neutrophils (Ly6G+) and macrophages (F4/80+) in Collagen-VI KO tumors. Thus, while cancer cell-derived Collagen-VI could be important for the formation of metastasis at multiple organ sites, other cell types in the metastatic microenvironment may be able also deposit it.

## Discussion

Pancreatic tumors are hallmarked by their stiff collagen-rich stroma. However, the metastatic cascade requires cells to adapt to softer environments, raising the question of how this plasticity is achieved. Culturing PDAC cells on hydrogels of varied stiffnesses showed substantial reprogramming of cells, which was accompanied by morphological changes that have been reported previously (Papalazarou et al., 2020). Despite having the same driver mutations (KRas^G12D^ and p53^R172H^), the two PDAC cell lines isolated from KPC mice showed unique responses to culture on soft vs stiff hydrogels, both at the individual gene expression and gene-program level. This reflects the heterogeneity of PDAC cell identity also observed in the clinical setting, that contributes to pancreatic cancer resistance to therapeutic intervention (Carstens et al., 2021). Given this, it is noteworthy that an upregulation of Collagen-VI, and matrisome genes more broadly, was a shared response of both PDAC lines to culture on soft substrate. ECM secretion by cancer cells in low-adhesion environments has been suggested previously, such as increased expression of SPARC, MPG and SPON2 in circulating tumor cells (CTCs) from patients (Ting et al., 2014). However, direct evidence for specific stiffness-dependent expression of ECM proteins by PDAC cells, as presented here, has been lacking. Our data adds to a growing repertoire of mechanosensitive mechanisms that PDAC cells use to survive and grow in different niches.

Our data disrupting various components of the mechanosensation pathway through plating cells on substrates that do not engage integrins, disruption of vinculin or talin interactions or knockdown of ILK (Figure 2) strongly suggests that increased Collagen-VI expression on soft substrates is due to a lack of integrin receptor engagement and downstream YAP nuclear translocation. How reduced YAP-driven transcription leads to increased Collagen-VI transcription is currently unclear. YAP coordinates expression of large gene networks, including those that can actively repress expression of other genes (Kim et al., 2015). Thus, increased Collagen-VI on soft substrata could be a consequence of relieved inhibition of another factor. Alternatively, inactivation of YAP may free up TEAD transcription factors to interact with other co-factor TFs such as VGLLs (Zhang et al., 2017) to promote expression. The differing responses of PDAC-A and PDAC-B cells to soft hydrogels highlights the complexity of mechanosensitive expression and suggest that exact pathways activated may differ depending on other properties or cell states.

An increasingly important question in relation to cancer ECM is the cellular origin and mechanism of action of specific ECM components. Along with most collagens, Collagen-VI expression is upregulated in primary PDAC and mostly derives from myofibroblasts (Tian et al., 2019). This is supported by data here showing that PDAC cells downregulate their own expression of Collagen-VI in stiff environment such as primary PDAC tumors. Stromal-derived Collagen-VI may still play important roles in PDAC progression, such as in promoting motility as has been reported for breast cancer (Wishart et al., 2020). Indeed, our data showing increased motility of PDAC cells on Collagen-VI coated coverslips supports this. Comparing metastatic tumors formed by PDAC cells with or without *Col6a1* KO, it is clear that cancer cells are a significant source of Collagen-VI in this context.

Our data shows that cancer cell-derived Collagen-VI has cell autonomous roles in determining metastatic potential of PDAC cells. Using a number of approaches, we show that Collagen-VI supports invasive potential of PDAC cells, *in vitro* and *in vivo*. This aligns with other recent studies reporting promotion of migration and invasion by Collagen-VI in other cancers (Chen et al., 2013; Wishart et al., 2020). Given that we see enhanced migration of PDAC cells on Collagen-VI coated coverslips, our data support a mechanism whereby motility effects are due to secreted Collagen-VI providing a local substrate for cancer cells. However, we cannot discount the possibility that Collagen-VI may have additional intracellular functions or that Collagen-VI may accumulate in cells on soft substrates due to inability to be secreted. Additionally, the COL6A3 fragment, endotrophin (ETP), has been suggested to enhance EMT in cancer cells (Park and Scherer, 2012), although it was not responsible for migration effects in breast cancer cells (Wishart et al., 2020). Finally, Collagen-VI may also support establishment of metastases by providing ECM peptides for attachment to the liver vasculature during early seeding, as has been shown for PDAC cell-derived matrisome proteins SERPINB5 and CSTB (Tian et al., 2020).

In summary, this study uncovers a mechanosensitive mechanism whereby pancreatic cancer cells alter their own extracellular matrix environment to support metastatic colonisation. Using a combination of gene expression profiling, *in vitro* characterisation and *in vivo* data we show how Collagen-VI expression is dependent on substrate stiffness. Our *in vivo* data also suggest that Collagen-VI is an important factor in the liver metastatic niche and can be provided either by the tumor cells or the liver resident cells. Our study highlights the dual nature of mechanosensation – as a response to soft as well as stiff environments – and illustrates the importance of the former to metastatic dissemination.

---

**Supplementary Figure 1.**
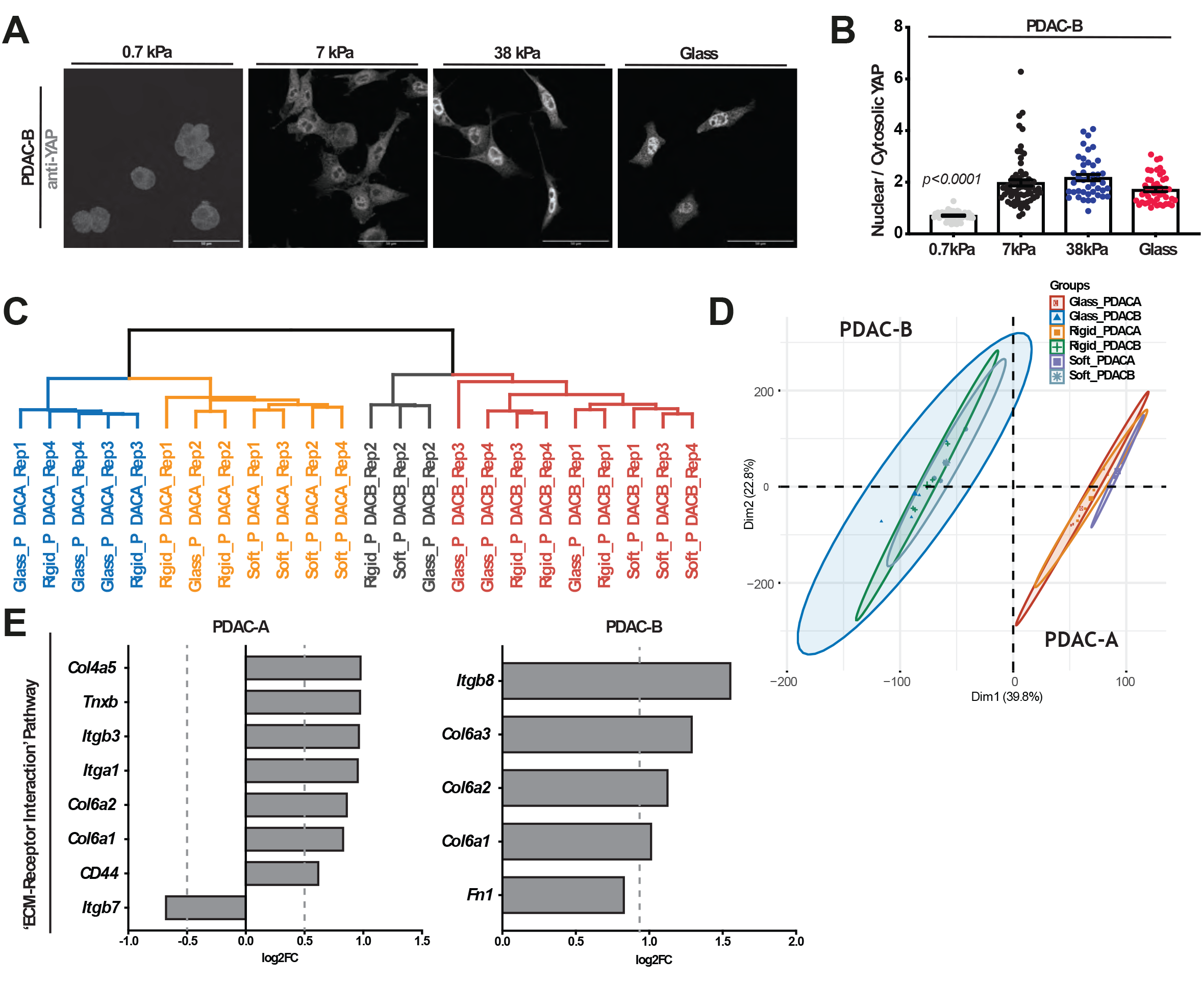
Low substrate stiffness alters the expression of matrisome-related genes in PDAC cells. **A**: Immunofluorescence of PDAC-B cells cultured atop of 0.7 kPa, 7 kPa or 38 kPa fibronectin-coated polyacrylamide hydrogels for 24 hours, showing YAP1 (grey). Scale bars, 50 μm. **B**: Quantification of ‘nuclear YAP1’/ ‘cytosolic YAP1’ intensity ratio for cells in (A). Values are mean ± s.e.m. from n= 52 cells, 0.7kPa; n= 68, 7kPa; n= 42, 38kPa; n= 46, glass. Cells are from 3 independent experiments. Statistical significance was assessed by Kruskal-Wallis test with Dunn’s multiple comparisons test. **C**: Hierarchical clustering of RNA sequencing signatures from PDAC-A and PDAC-B cells cultured on fibronectin-coated 0.7 kPa, 38 PAAm hydrogels and glass coverslips for 24 hours. **D**: PCA clustering of cells from (C). **E**: Bar plot displaying differentially expressed genes (p_adj_ < 0.05; log2 fold change >1) from the ‘ECM-Receptor Interaction’ KEGG pathway that were enriched in cells from (C). Data is organised by log2 fold change.

**Supplementary Figure 2.**
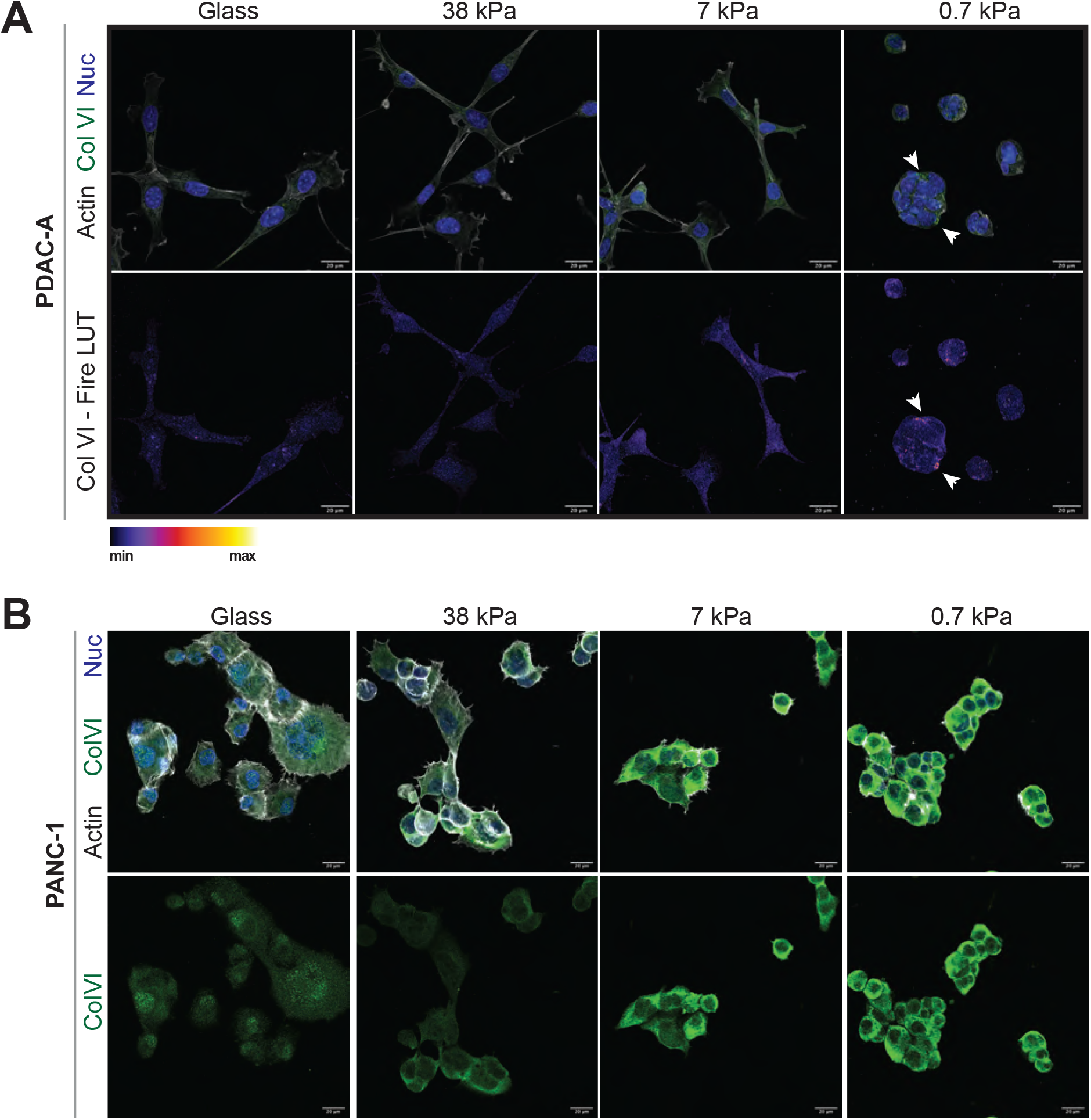
Collagen VI is upregulated in PDAC cells upon low substrate stiffness. **A**: Top; Immunofluorescence of PDAC-A cells cultured on glass coverslips, 0.7-, 7- and 38-kPa fibronectin-coated hydrogels, showing Collagen VI (green), Actin (grey) and nuclei (blue). Representative pictures from 3 independent experiments. Bottom; Individual Collagen VI channel (Fire LUT). Scale bars, 20μm. Arrowheads indicate Collagen VI enrichment. **B**: Top; Immunofluorescence of PANC-1 cells cultured on glass coverslips, 0.7-, 7- and 38-kPa fibronectin-coated hydrogels, showing Collagen VI (green), Actin (grey) and nuclei (blue). Representative pictures from 3 independent experiments. Bottom; Individual Collagen VI channel (Green). Scale bars, 20μm.

**Supplementary Figure 3.**
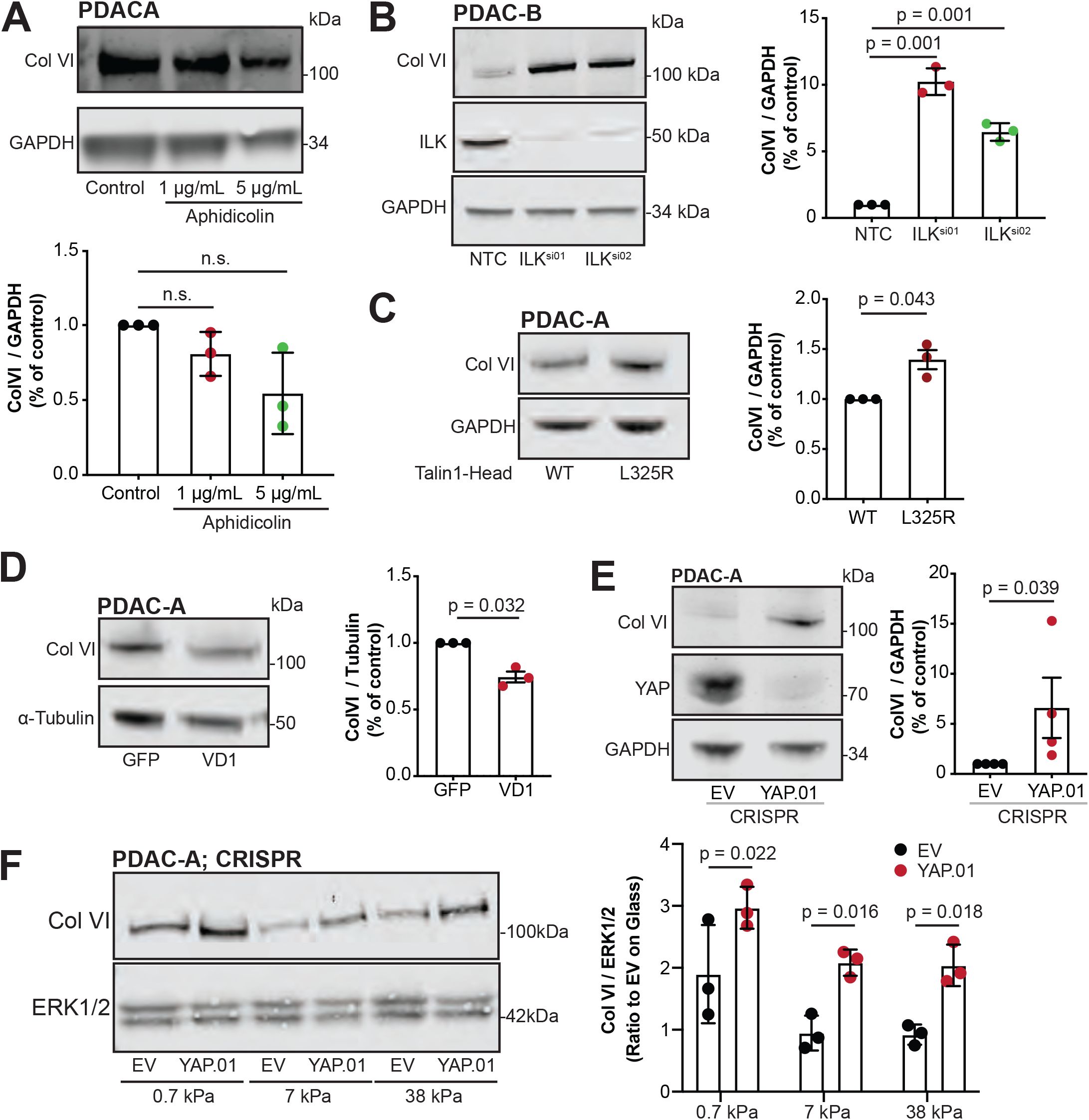
Loss of ECM adhesion and mechanosensing upregulates Collagen VI expression in PDAC cells. **A**: Top; PDAC-A cells were treated with 1 μg/mL and 5 μg/mL aphidicolin for 24 hours and immunoblotted for Collagen VI and GAPDH (loading control). Pictures are representative of 3 independent experiments. Bottom; Densitometric quantification of protein expression. Values are mean ± s.d. B: Left; Control (NTC) or ILK silenced (*Ilk*^si01^, *Ilk*^si02^) PDAC-B cells were immunoblotted for Collagen VI, ILK and α-Tubulin (loading control). Right; Densitometric quantification of protein expression. Values are mean ± s.d. **C**: Left; PDAC-A cells expressing either Talin 1-head domain (WT, control) or Talin-head L325R mutant (L325R) were immunoblotted for Collagen VI and GAPDH (loading control). Right; Densitometric quantification of protein expression. Values are mean ± s.d. **D**: Left; PDAC-A cells expressing either GFP (control) or GFP-tagged Vinculin Domain 1 (VD1) were immunoblotted for Collagen VI and α-Tubulin (loading control). Blots are representative of three independent experiments. Right; Densitometric quantification of protein in C. Values are mean ± s.d. E: Left; Control (EV) or YAP-depleted (YAP.01) PDAC-A cells were immunoblotted for Collagen VI, YAP and ERK1/2 (loading control). Pictures are representative of 4 independent experiments. Right; Densitometric quantification of ColVI protein. Values are mean ± s.d. **F**: Left; Control (EV) or YAP-depleted (YAP.01) PDAC-A cells were cultured on fibronectin-coated 0.7-, 7- and 38-kPa hydrogels and were immunoblotted for Collagen VI and ERK1/2 (loading control). Right; Densitometric quantification of protein. Values are mean ± s.d. and representative from 3 independent experiments. Statistical significance was assessed by two-way ANOVA and p-values were corrected for multiple comparisons by Šídák’s test. All data in A-D are from 3 independent experiments. Statistical significance was assessed by two-tailed one-sample *t*-test on natural log-transformed values.

**Supplementary Figure 4.**
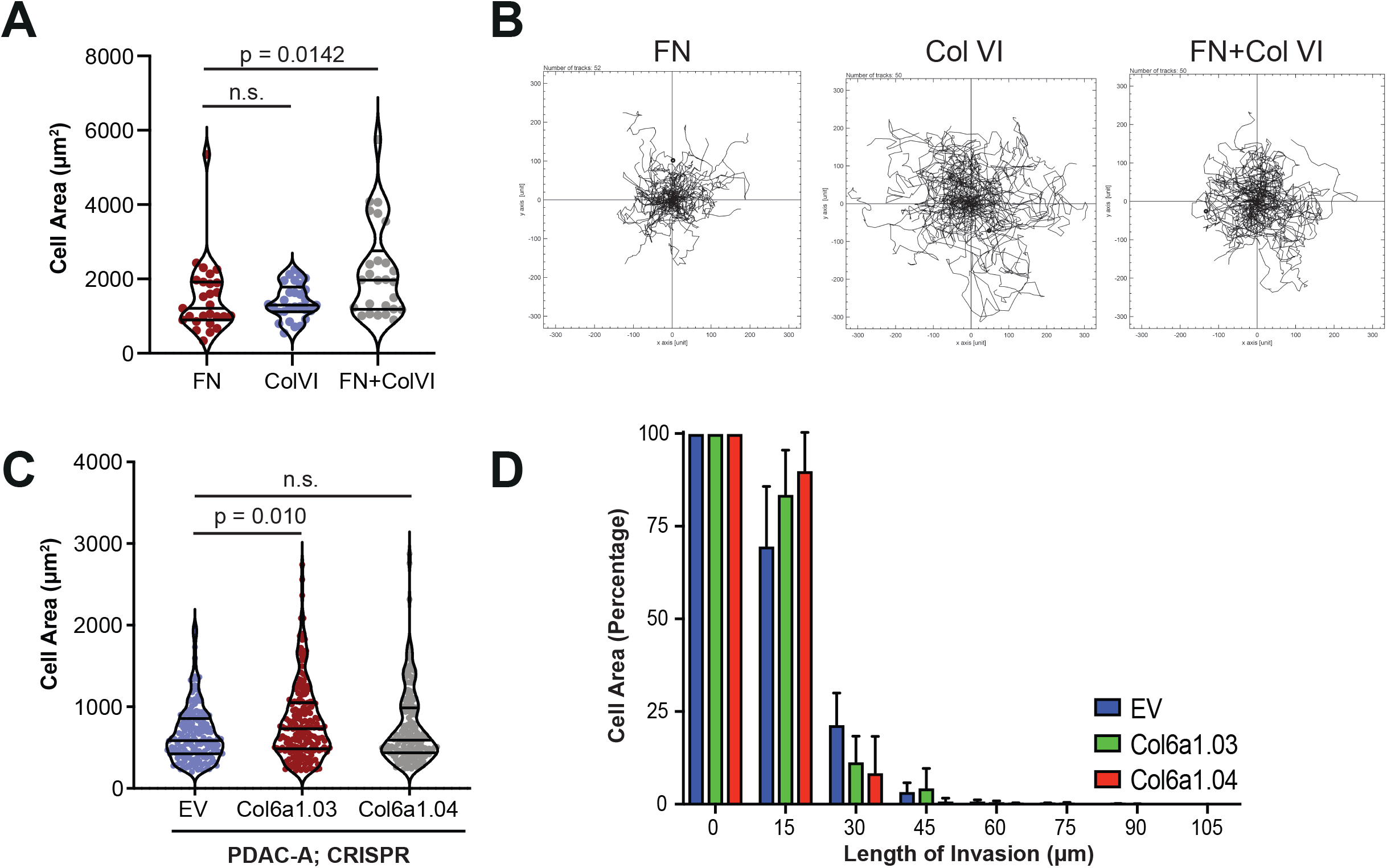
Collagen VI ECM supports migratory behaviour of PDAC cells *in vitro* and loss of Col6a1 expression delays invasion through recombinant basement membrane ECM. **A**: Cell area (μm^2^) quantification of PDAC-A cells cultured on fibronectin (FN), collagen VI (ColVI) or fibronectin and collagen VI (FN+ColVI) glass coverslips. Values are from *n* = 27 FN cells; *n* = 29, ColVI; *n* = 26, Fn+ColVI cells. Cells are from three independent experiments. Statistical significance was assessed by Kruskal-Wallis with Dunn’s multiple comparisons test. **B**: Tracks (spider plots) of PDAC-A cells migrating on fibronectin (FN), collagen VI (ColVI) or fibronectin and collagen VI (FN+ColVI) glass coverslips for 16 hours. **C**: Cell area (μm^2^) quantification of Control (EV) or Collagen VI depleted (Col6a1.03 and Col6a1.04) mouse PDAC-A cells. Values are mean ± s.d. from n=170 EV, n=191 Col6a1.03 and n=183 Col6a1.04 cells from 3 independent experiments. Statistical significance was assessed by Kruskal-Wallis with Dunn’s multiple comparisons test. **D**: Quantification of invaded area of Control (EV) or Collagen VI depleted (Col6a1.03 and Col6a1.04) mouse PDAC-A cells invading through the inverted invasion assay setup. Intensity for each depth is reported as a percentage of intensity at 0 μm. Values are mean ± s.d. from 3 independent experiments.

**Supplementary Figure 5.**
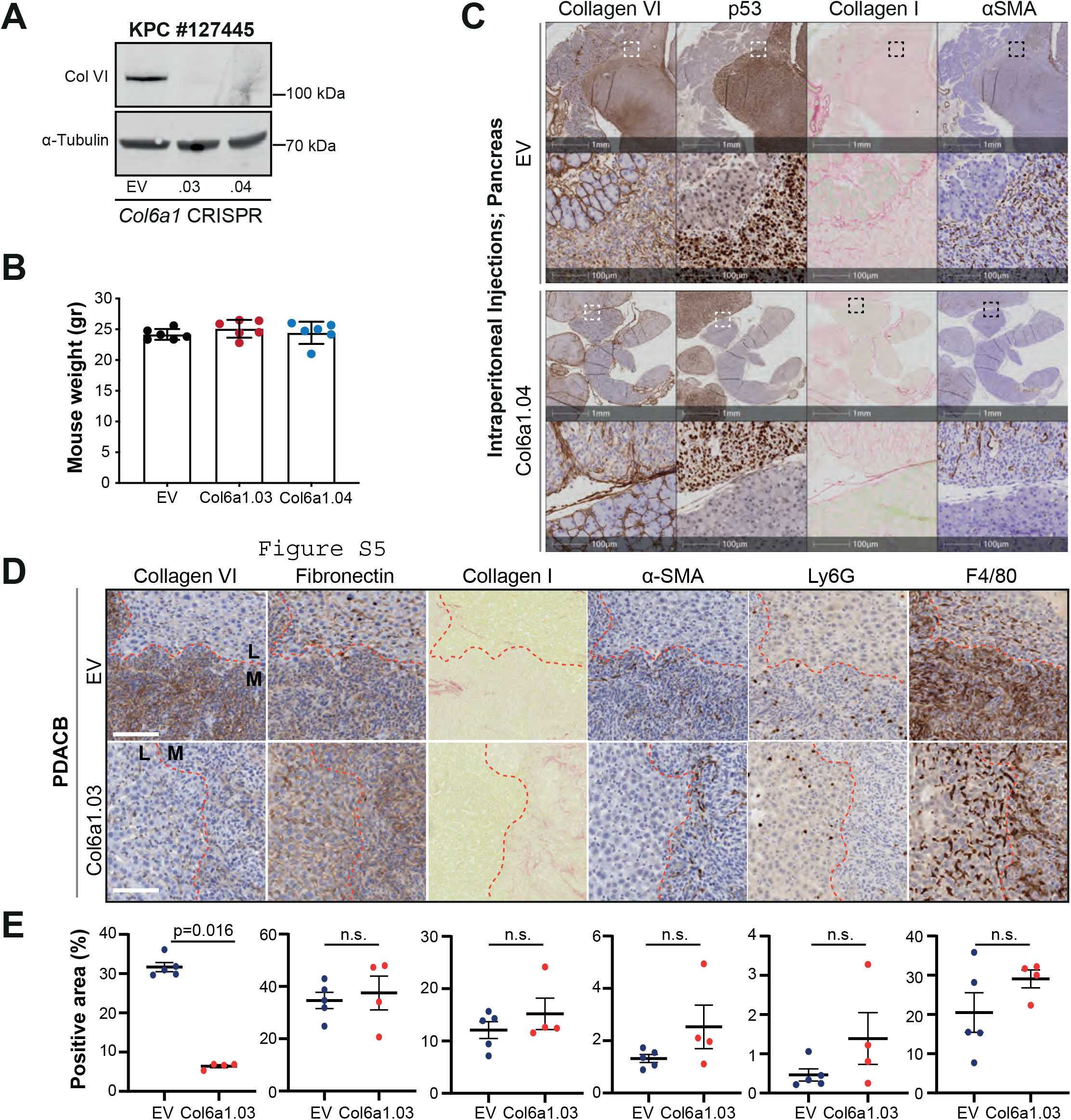
Collagen VI expression supports establishment of pancreatic metastasis in vivo. **A**: Control (EV) or Collagen VI depleted (Col6a1.01-04) KPC cells were immunoblotted for Collagen VI and *α*-Tubulin (loading control). **B**: Weight (gr) per mouse as indicated from intraperitoneal injection of control (EV) or Collagen VI depleted (Col6a1.03 and Col6a1.04) KPC cells after sacrifice. Values are mean ± SD from n = 6 EV, n = 6 Col6a1.03 and n = 6 Col6a1.04 mice. **C**: Representative immunohistochemistry images showing Collagen VI, p53, Collagen I and α-SMA expression in tumors formed in the pancreas by intraperitoneal injection of control (EV) (top 2 panels) or Collagen VI depleted (bottom 2 panels) KPC cells. Scale bars, 1mm and 100μm. **D**: Representative immunohistochemistry images showing Collagen VI, Fibronectin, Collagen I, α-SMA, Ly6G and F4/80 expression in liver metastatic nodules formed by intrasplenic injection of control (EV; top) or Collagen VI depleted (Col6a1.03; bottom) KPC cells. Red line denotes liver (L) and metastasis (M) boundary. Scale bars, 100μm. **E**: Quantification of positively stained regions over tumor area (%) from D. Values are mean ± s.e.m. from n = 5 control (EV) and n = 4 Col6a1.03 mice. Statistical significance was assessed by Mann-Whitney test (Collagen VI) and unpaired t-test (Fibronectin, Collagen I, αSMA, Ly6G and F4/80).

